# Evaluating sustainable feeds for aquaculture by simulating Atlantic salmon lipid metabolism

**DOI:** 10.1101/2024.06.01.596980

**Authors:** Filip Rotnes, Jon Olav Vik, Ove Øyås

**Affiliations:** Faculty of Chemistry, Biotechnology and Food Science, Norwegian University of Life Sciences (NMBU), Christian Magnus Falsens vei 18, Ås, 1433, Norway; Faculty of Biosciences, NMBU, Oluf Thesens vei 6, Ås, 1433, Norway

**Keywords:** Atlantic salmon, sustainable feeds, metabolic model

## Abstract

Atlantic salmon aquaculture is an important food source globally, but its sustainability is challenged by environmental impacts and the nutritional demands of farmed fish, particularly when it comes to fatty acids. Salmon feeds still rely heavily on fish or soybean meal, which poses sustainability concerns due to overfishing and carbon footprint. Innovations in feed composition seek to address these challenges, e.g., by using more sustainable ingredients, but the impacts of alternative feeds on fish and environment can be hard to quantify. Here, we developed a model with detailed and flexible accounting for lipids – Simulated Salmon Lipid Metabolism (SimSaLipiM) – to predict the nutritional and environmental outcomes of feed formulations. Integrating SimSaLipiM with feed ingredient databases enabled detailed analysis of an *in vivo* feed trial *in silico*. The model predicted optimal feed efficiency in agreement with observations as well as a detailed energy budget and fish biomass lipid composition for each feed. We also used the model to formulate novel sustainable feeds and feed supplements by minimising CO_2_ footprint. Thus, SimSaLipiM makes it easy to identify recipes that optimize key feed properties such as efficiency and environmental impacts. This could be a valuable tool for feed manufacturers, guiding the formulation of feeds that are both sustainable and cost effective. By bridging the gap between feed formulation and the flexible growth and energy requirements of a fish, SimSaLipiM can contribute to advancing sustainable aquaculture.

## 1. Introduction

Atlantic salmon farming generates great value [1] but faces challenges of sustainability and nutritional value [2, 3]. Wild salmon are carnivores, but scarcity of fish-based feeds has driven a shift to plant-based sources of fat and protein in salmon farming. Although a remarkable feat of feed and nutrition technology, this shift affects salmon health and the nutritional value of the product [3]. Plant oils have a higher ratio of omega-6 (*ω*-6) to omega-3 (*ω*-3) than marine oils, reducing the robustness of farmed salmon to stress [4] and diminishing the nutritional benefit of the meat as a dietary polyunsaturated fatty acid (PUFA) source [5]. Soybean protein content is also associated with gut inflammation, affecting fish growth [6]. Moreover, the feedstuff market can be highly volatile, with pricing and availability of feed ingredients shifting rapidly. The environmental footprint is substantial even for plant-based feeds [2], and new ingredients such as kelp, microbial meal, or insect meal are being investigated as alternative sources of beneficial fatty acids and amino acids [7]. Thus, a key challenge is to compose serviceable feeds that balance economic profit with fish health, the nutritional value of salmon meat, and an acceptable environmental footprint.

The SALARECON metabolic model for Atlantic salmon has brought the power of constraint-based modelling to production biology [8], potentially opening new opportunities for efficient and sustainable meat production [9]. SALARECON has already been used to explain salmon growth and predict amino acid supplements for improved utilisation of commercial feeds [8], to predict combinations of commercial feeds that maximise profit [10], and to functionally interpret gene expression data [11]. However, its rudimentary representation of lipids includes only synthesis up to palmitate (lipid number C16:0), and it does not cover the variety of long-chain and unsaturated fatty acids found in available feed ingredients nor the diversity of fatty acyl glycerides and phospholipids found in salmon tissue. This represents a major limitation, as the fatty acid composition of salmon tissues, which is largely determined by the composition of the feed, is of primary importance both for the fish and for the nutritional value of its meat [4, 5].

In a metabolic model, growth and associated energy costs are represented by a biomass reaction that traditionally has a rigid ratio between building block and energy molecules [12], but biomass compositon is known to be condition-dependent in general [13, 14] and for salmon lipids in particular [15, 16]. Also, simplified representation of biomass lipids is common in metabolic models due to the combinatorial complexity of lipid metabolism [17]. However, progress has been made through semi-automated reconstruction of lipid metabolism, most notably for yeast [18, 19], as well as large manual reconstruction efforts, for example for humans [20, 21] and other animals through orthology [22]. Still, despite lipids being major constituents and energy sources of fish and many similarities between fish and human lipid metabolism [23], none of the available manually curated models of fish account for lipid metabolism in any detail [8, 24]. Doing so would enable more accurate simulated feed trials *in silico* that could help predict novel sustainable feed compositions for aquaculture *in vivo*.

In this study, we present a new metabolic model for Atlantic salmon – Simulated Salmon Lipid Metabolism (SimSaLipiM) – with detailed and empirically based fatty acid metabolism. Specifically, we extended SALARECON with a lipid metabolism module accounting for synthesis, elongation, desaturation, and beta oxidation of long-chain fatty acids in Atlantic salmon. Given a list of fatty acids to biosynthesize and degrade, this module generates the SimSaLipiM model, which also integrates measured fatty acid compositions from salmon to allow for a flexible and unbiased biomass composition. Feed supply reactions provide metabolites at rates proportional to the observed molecular composition of feed ingredients and allow direct prediction and simulation of feed compositions. Thus, the model accounts for a flexible lipid composition in the filet within biologically observed boundaries, and it allows for *in silico* feed trials, which we performed in three ways: (1) to validate the model and interpret data from an *in vivo* feed trial mechanistically, (2) to identify novel feed recipes with minimal CO_2_ footprint, and (3) to predict beneficial ingredient additions to improve feed performance. Our results demonstrate that SimSaLipiM can help guide development of new feeds for aquaculture that are efficient as well as sustainable, potentially reducing the need for expensive *in vitro* and *in vivo* experiments.

## 2. Methods

### 2.1. Extending models with fatty acid metabolism

A general framework was made to add fatty acid metabolism to a constraint-based model based on user input. The fatty acids to include are defined in a table, in which each fatty acid is identified by a lipid number and additional information such as metabolite IDs, names for the fatty acid in its free form and when bound to coenzyme A (CoA), and a KEGG compound ID for the fatty acyl-CoA. The chemical formula of each molecule is inferred from the lipid number, and KEGG IDs can be used to search for further annotation. Metabolite identifiers for free and CoA-bound fatty acids are based on the lipid number and cellular compartment unless other identifiers are stated. For example, the fatty acyl-CoA with lipid number C22:1n9 in the cytoplasm will be named “c221n9coa_c” if no other identifier is specified.

Fatty acid reactions are grouped into elongation, desaturation, and beta oxidation. Desaturation is a one-reaction process, while elongation and beta oxidation are multi-step processes with intermediate metabolites. Intermediates are assigned identifiers based on the shortest fatty acid in the process, plus an indication of modifications such as “oh” for hydroxyl, “cooh” for carboxyl, and so on. Elongation, desaturation, and beta oxidation reactions are defined in tables, in which each reaction type is assigned an ID prefix, a reaction name, a compartment in which the reaction occurs, EC numbers, cofactors, byproducts, genes, and a substrate length. Additionally, beta oxidation reactions are characterised by their specificity for double bond positions on the substrate. For each fatty acid to include in the model, the possible reaction processes are identified and then evaluated based on their predicted products. Elongation and desaturation reactions are only added if the product of the process is in the table of target fatty acids. Beta oxidation is added for all fatty acids, however, as all fatty acids are assumed to be digestible. Identifiers for reactions are generated in a systematic manner by combining the reaction’s prefix with the identifier of the end product of the process. For instance, the end product of one beta oxidation process of c22:1n9 would be c20:1n9.

### 2.2. Simulating feeds through feed supply reactions

Feed supply reactions simulate ingestion of feed. Most feed ingredients come from biological sources and are, in essence, mixtures of chemical compounds. Design of diets is a venture towards ideal mixtures of compounds through combining available ingredients while keeping account of their cost. By using the amount of each compound (in mmol) per gram fed ingredient as stoichiometric coefficients in a feed supply reaction, we can simulate uptake of feed. The equation of the resulting pseudoreaction is empty on one side and has extracellular metabolites on the other side. This is similar to the conventional biomass reaction, but instead of consuming metabolites from the system, it supplies them. Superfluous metabolites and metabolic byproducts are excreted back to the environment via exchange reactions, which can be interpreted as a simulated fecal composition. Feed supply reactions replace the inward direction of exchange reactions, except for a few compounds such as oxygen. By adding several feed supply reactions, we can evaluate combinations of ingredients. When the model is optimised with flux balance analysis (FBA) [25], feed composition is determined by the choice of objective function. The ingredients can be valued equally or weighted by economic value, climate footprint, or other factors.

### 2.3. Building the SimSaLipiM model by extending SALARECON

We curated three tables specifying fatty acids to add to SALARECON [8] and the reactions that may connect them. One table contains the fatty acids to include, one describes the elongation and desaturation reactions, and one describes the beta oxidation reactions. Fatty acids were chosen based on those listed in the Feedtables database [26] and in whole-body composition data for Atlantic salmon from Tocher et al. [16] and Mock et al. [27]. Common saturated and desaturated fatty acids with even-numbered chain lengths up to 28 carbons were included, and among these are the desirable *ω*-3 fatty acids eicosapentanoeic acid (EPA) and docosahexanoeic acid (DHA). If available, a metabolite identifier from the BiGG database [28] was chosen, both for the free fatty acid and for the fatty acyl-CoA. The free and CoA-bound forms of the desired fatty acids were added to the model along with the reactions needed to interconvert them. Products of all elongation and desaturation processes were determined for all fatty acids, and if the product was also among the fatty acids in the model, the process was added. Beta oxidation reactions were added to break down all fatty acids.

We also added 321 feed supply reactions to the model based on molecular compositions from Feedtables in three steps: First, compounds listed in Feedtables were mapped to the namespace of the model. Second, this mapping was used to convert the listed mass fractions from gram per gram to mmol per gram using the molecular weight of the corresponding metabolites in the model. Finally, these values were used as stoichiometric coefficients in feed supply reactions. The result is one extracellular reaction per ingredient, with all metabolites on the product side. Metabolites are thus supplied in the same ratio as they are present in the feed ingredients used and may either be imported via transport reactions or excreted via exchange reactions.

### 2.4. Feed trial simulation and validation

Data from an experiment by Mundheim et al. [29] were used to simulate feed trials *in silico* for eight diet recipes designed to evaluate combinations of plant-based ingredients with fish meal of different qualities *in vivo*. Measurements of macronutrients in feed and fish were reported along with growth rates and feed efficiency measurements. The ingredients in the diet recipes were manually interpreted as ingredients listed in the Feedtables database, giving rise to a molecular composition of each feed. To account for differences between the experimental diets and the molecular interpretations, the simulated diets were uniformly scaled to the same dry matter content as the reference diets. Each resulting diet was used to define a feed supply reaction, and the growth rate of the model was fixed to the measured dry matter content of fish that were fed that diet, which is the same as 1 g/g wet weight. Using minimisation of the feed supply reaction as objective, parsimonious FBA (pFBA) [30] was used to find the minimal feed uptake while minimising total flux. The flux through the feed supply reaction can then be interpreted as a prediction of the feed conversion ratio (FCR), which was compared to FCRs reported by Mundheim et al. [29]. The predicted feed uptake was fixed and then the rate of ATP rephosphorylation was maximised. To interpret pFBA predictions as an energy budget, the energy content for any protein or lipid was assumed to be 17 or 37 kJ per gram, respectively [31] . These assumptions were used to calculate the energy content of biomass, faeces and feed. Compounds that were not characterised as fatty acids or amino acids were not assigned an energy content. ATP production was interpreted as activity, using 45 kJ per mol rephosphorylated ATP [31] . Energy of excretion of urea was calculated based on rate, formula weight, and the energy content of urea listed in Feedtables. Finally, energy loss to heat was calculated as the remainder when subtracting the other energy factors from the total energy content of feeds.

### 2.5. Predicting sustainable feeds

The ReCiPe database of the Global Feed LCA institute (GFLI) [32] contains 1499 measurements of ingredients with data for up to 21 life cycle analysis (LCA) categories. These can be used to characterise feed compositions in terms of environmental costs and as constraints in optimisation. Many of the rows also contain country of origin. To pair this with molecular compositions of ingredients, the ingredient names were mapped by the following algorithm: For each name in feedtables, the ReCiPe dataset was filtered by identifying the subset of ingredient names containing the first word of the feedtables ingredient. The resulting table was then filtered in the same manner using the second word, and so on until all words were matched or the search could not be made more specific. Precautions were taken to avoid some obvious errors. Maize and corn were added to a dictionary of synonyms, and made substitutable in the search. Aditionally, some entries in Feedtables seem to have the most characteristic word at the end of the name. Two types of entries were identified, namely entries starting with “Processed animal protein” or “Fish oil”. These were renamed with the species name as the first name, so that “Processed animal protein, pig” becomes “pig Processed animal protein, pig” and so on. After mapping rows of LCA data to molecular compositions of ingredients, 165 out of 321 feed ingredients had an associated set of measurements. For location-specific analysis, we filtered ingredient measurements by country in order to conserve the quality of the matching between local and global approaches. For both local and global values, the median value of CO_2_ emissions (including eutrophication) was used to define the objective of minimisation.

## 3. Results

As summarised in Fig. 1, we generated a model capable of simulating biosynthesis and degradation of 49 fatty acids with a flexible biomass composition based on Atlantic salmon whole-body composition data [15, 16]. The flexible biomass composition is achieved by distributing the biomass reaction across separate reactions that produce the individual biomass components: protein, lipids, carbohydrates, RNA, and DNA. To account for the observed flexibility of the lipid composition, we used the collected whole-body measurements to formulate 58 feasible biomass lipid compositions covering a range of conditions. Any linear combination of these empirical fatty acid compositions is available to the model, as illustrated by the allowed ranges of biomass lipid stoichiometric coefficients and corresponding ratios between *ω*-3, *ω*-6, and other fatty acids. The SimSaLipiM model generated in this paper represents Atlantic salmon genome-scale metabolism as 1252 genes encoding enzymes that catalyze 1975 reactions interconverting 1642 metabolites. We validated the model by evaluating its ability to perform a range of metabolic tasks [21, 33]. SimSaLipiM was able to perform a much larger share of tasks than SALARECON in lipid metabolism [8], reflecting the expansion of scope with emphasis on fatty acid metabolism. We also used Memote, a standardised tool for model testing, to evaluate the consistency and annotation of the model, achieving a score of 92%(maximum score is 100%).

**Figure 1:**
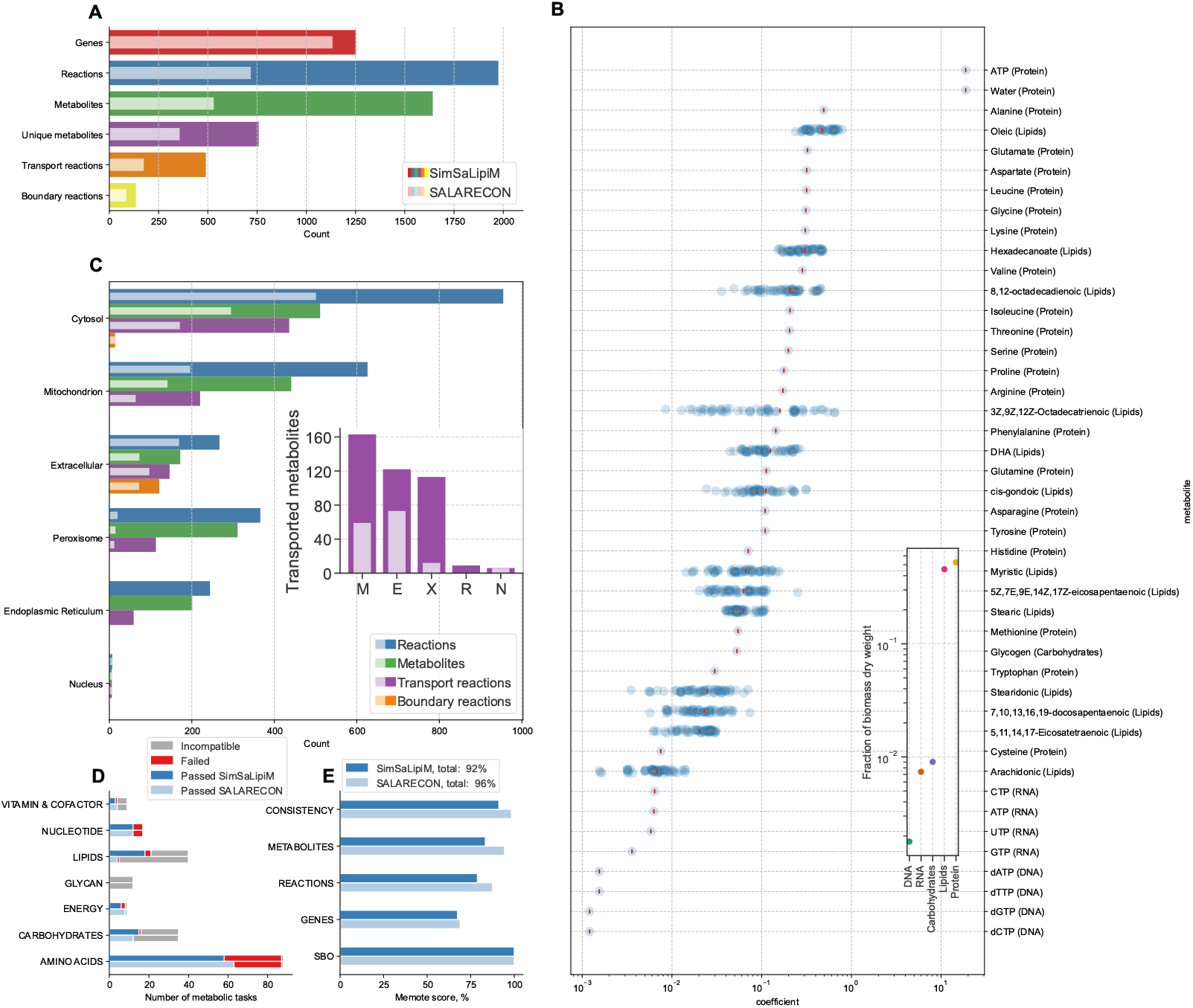
Contents and predictions of SimSaLipiM compared to SALARECON. (**A**) Number of genes, reactions, and metabolites in SimSaLipiM and SALARECON. The number of unique metabolites is also shown because the same metabolite can be found in multiple compartments. Reactions include transport reactions, which transport metabolites between compartments, and boundary reactions, which allow metabolite exchange with the environment. (**B**) Stoichiometric coefficients of metabolites in the biomass reaction. The composition of the lipid fraction of the biomass was made flexible by using empirical fatty acid ratios to formulate multiple pseudoreactions for lipid biosynthesis. There are therefore multiple biomass coefficients for fatty acids. The inset shows the fraction of biomass dry weight assigned to protein, lipids, carbohydrates, RNA, and DNA. (**C**) Metabolites and reactions are distributed across six compartments for SimSaLipiM and SALARECON. Transport reactions are associated with two compartments. The inset displays the number of metabolites transported in or out of the mitochondrion (M), extracellular space (E), peroxisome (X), endoplasmic reticulum (R), and nucleus (N). (**D**) Metabolic task performance for SimSaLipiM and SALARECON grouped by metabolic subsystem. Each column pair illustrates the number of tasks that succeed, fail, or are incompatible with SimSaLipiM or SALARECON, respectively. (**E**) Model quality scores from MEMOTE for SimSaLipiM and SALARECON.

Fig. 2 summarises the use of molecular compositions from the Feedtables database to convert diet recipes from an *in vivo* feed trial [29] into feed supply reactions. The recipes were formulated with varying amounts of marine and plant-based ingredients to compare high- and low-quality fish meal. Specifically, recipes 1–4 contained high-quality fish meal and recipes 5–8 contained medium-quality fish meal. As expected, we found clearest separation between recipes using high- and medium-quality ingredients and grouping of recipes containing the same ingredients. Putative molecular compositions were obtained by manually mapping the ingredients in the diets to entries in the Feedtables database. The resulting molecular diets were then scaled to contain the same amount of metabolites as the reference diets in terms of mass, and we found that the molecular diets clustered based on fish-to-plant ingredient ratio.

**Figure 2:**
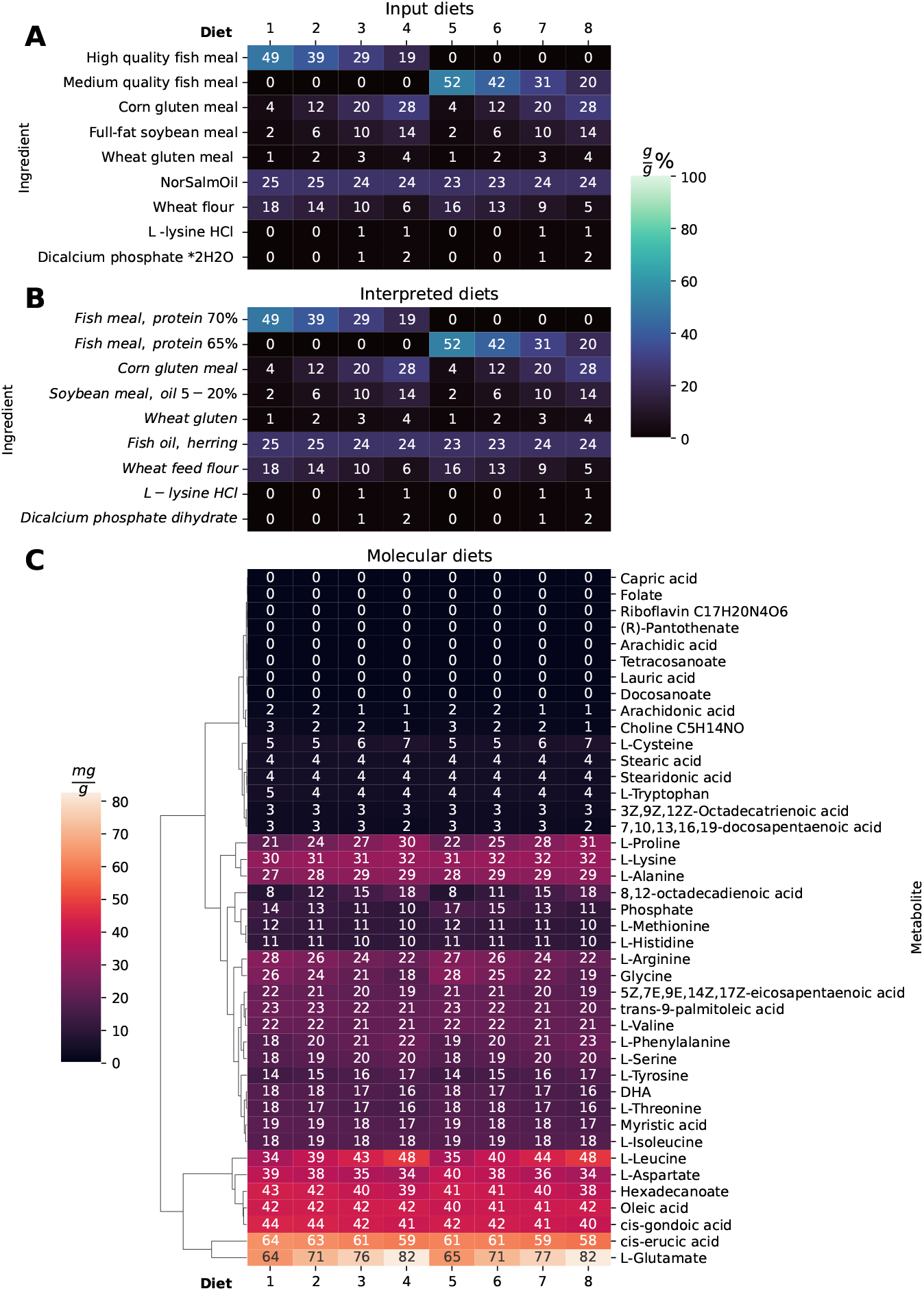
Translating feed recipes to metabolites via the Feedtables database. (**A**) Recipes from an *in vivo* feed trial [29] where fish meal of high (diets 1–4) and medium (diets 5–8) quality was combined with plant-based ingredients in different ratios. The ratio of plant-based to marine ingredients increased from 1 to 4 and from 5 to 8. (**B**) Recipes after manual mapping of feed ingredients to the Feedtables database. (**C**) Molecular compositions of diets after mapping to Feedtables. Diets were scaled to contain the same mass of metabolites as the reported diet dry weight without ash and fibre.

As shown in Fig. 3, recipes and body composition in terms of macromolecules from the *in vivo* feed trial [29] were used to perform an *in silico* feed trial, predicting FCR as well as corresponding energy budgets and salmon biomass fatty acid compositions. Total energy available in the feed was decomposed into a hierarchy in which dietary energy can leave the system through faeces, urea, heat, growth, or maintenance/activity. All optimal FCR predictions were slightly below but close to the reference values, while FCRs for some of the individual biomass lipid compositions were up to five times higher than the reference values, highlighting the importance of a flexible representation of biomass lipids for accurate predictions. The variation in predicted energy budget and FCR between diets was also substantial but comparable within low- and high-quality diet pairs with the same ratio of fish- to plant-based ingredients. As expected, the amount of low-quality feed needed per gram growth was predicted to be higher than that of high-quality feed, both in terms of total energy and in terms of energy that could not be utilised for growth. With the exception of diets 2 and 5, the predicted total amount of energy expended was higher for low-quality than for high-quality diets across categories in the simulated trial.

**Figure 3:**
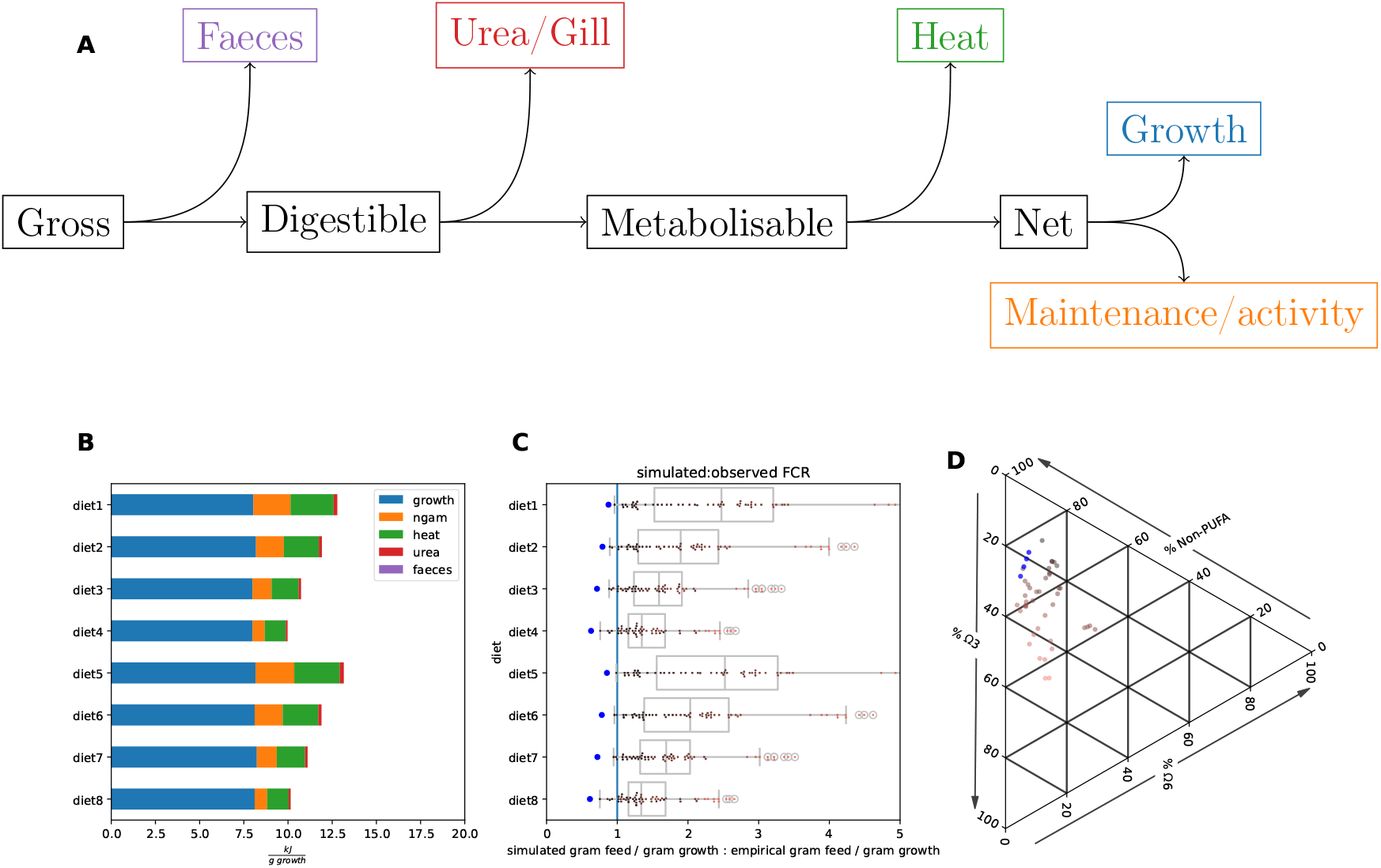
Predicting energy budget, feed efficiency, and biomass fatty acid composition for an *in vivo* feed trial. (**A**) Decomposing feed energy contents. The gross energy in the feed is broken down to digestible energy and energy loss as faeces. Digestible energy is then converted to metabolisable energy with energy loss as urea, and the metabolisable energy can be broken down to net energy with energy loss as heat. Net energy can be allocated to growth or maintenance and activity. (**B**) Predicted energy budget for diets from an *in vivo* feed trial [29] using maximum utilisation of the diet as objective. Diets 1–4 and 5–8 consist of the same ingredients with different ingredient ratios. (**C**) Predicted FCR relative to experimental FCR. The blue dot is the optimal value that minimises FCR with a flexible fatty acid biomass composition, which corresponds to the energy budget in B. The other dots are predicted FCRs for each invididual empirical fatty acid composition allowed in the biomass reaction colored by biomass *ω*-3 content. (**D**) Fatty acid biomass compositions corresponding to the energy budget in B and the minimal FCR in C. The proportional composition of *ω*-3, *ω*-6, and non-PUFA in biomass are shown in a ternary diagram.

Optimal feed composition is dependent on the objective used for optimisation, which is itself influenced by several factors such as time and place or assumptions of optimality based on evolution and selective breeding. Fig. 4 illustrates optimisation of feeds under different circumstances and using different objective functions. First, optimal feed recipes were determined for each of the 58 fatty acid compositions in the model. Specifically, minimal feed recipes were predicted for each of the empirical fatty acid compositions along with the corresponding biomass fatty acid composition. Using body *ω*-3 contents in terms of mass as metric, the optimal combination of fatty acid-rich ingredients in feed was identified to be a mixture of sardine oil and sunflower oil for high-*ω*-3 body compositions. Second, we used CO_2_ footprint as the objective function, based on data from Feedtables and ReCiPe and performing simulations for four different fixed FCRs to avoid inordinate amounts of ingredients with miniscule carbon footprint. Predicted fatty acid compositions showed little variation in PUFA content and *ω*-3-to-*ω*-6 ratios beetween databases and FCR levels, but *ω*-3 levels were higher for ReCiPe than for Feedtables. Finally, by filtering ingredients by country in the ReCiPe dataset, optimal feeds were identified for the major aquaculture producers Canada, Chile, Norway, UK, and USA. Both the CO_2_ footprints for the same ingredient and the availability of data on ingredients differed between countries. Concentrated ingredients such as potato protein extract, pea protein extract, soybean protein concentrate, wheat gluten, and oils from plants and fish were among the top predicted ingredients.

**Figure 4:**
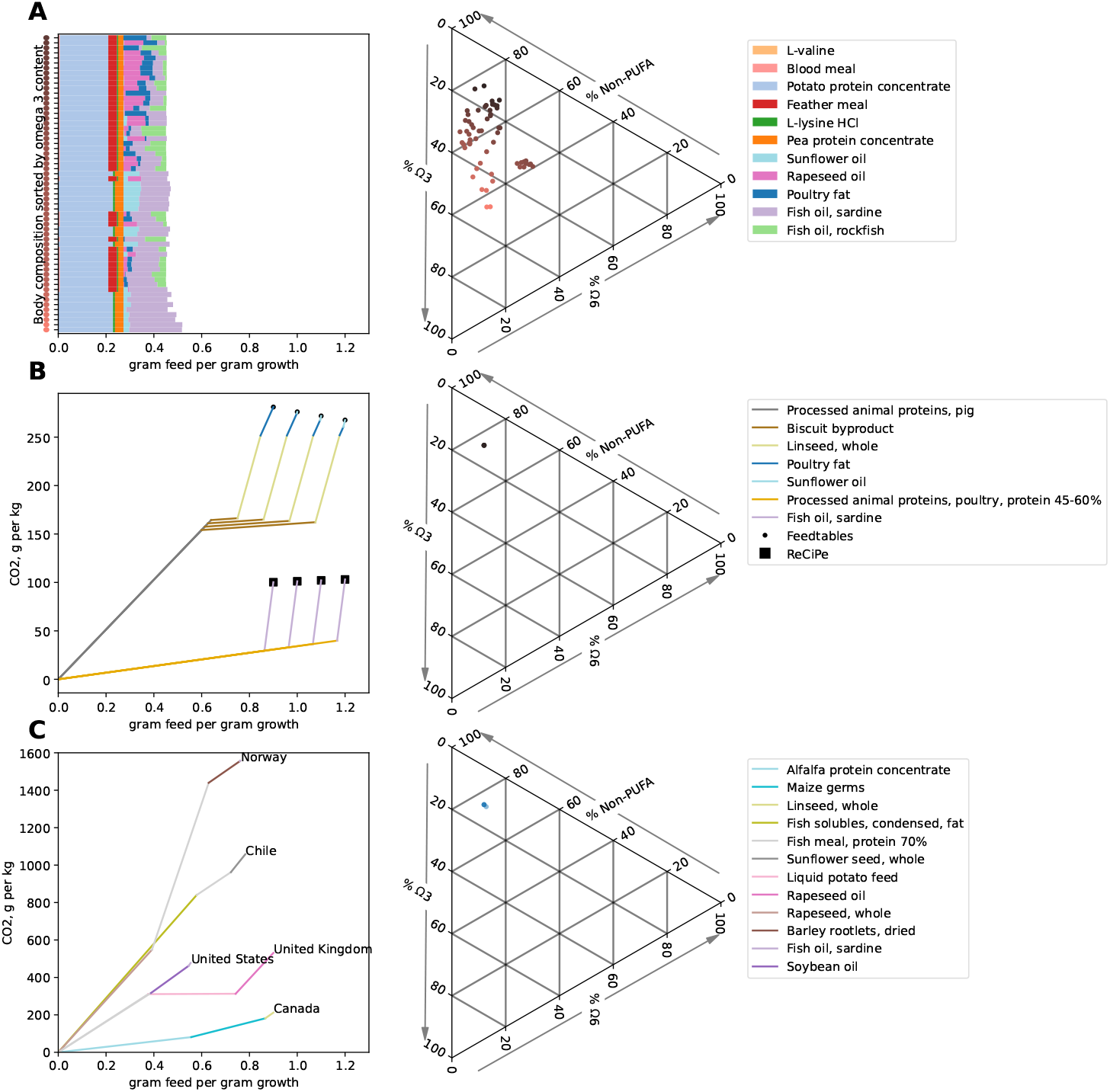
Predicting new feeds that optimise efficiency and CO_2_ footprint. (**A**) Feed composition that minimises FCR (grams feed per gram growth) for each empirical biomass fatty acid composition in the model. The left side shows FCR with optimal compositions sorted from highest to lowest biomass *ω*-3 content. The predicted optimal recipes vary in the lipid-rich ingredients, combining rockfish oil, lard, poultry fat, and sardine oil. The right side shows the proportion of *ω*-3, *ω*-6, and non-PUFA in the empirical biomass compositions. (**B**) CO_2_ emissions at four different feed conversion ratios for predicted feeds that minimise CO_2_ footprint (grams CO_2_ per gram growth) based on LCA data for ingredients found both in Feedtables (circles) and ReCiPe (squares). Each line represents addition of an ingredient to the diet, ingredients are added in order of increasing contribution to the CO_2_ footprint, and colors identify each ingredient. (**C**) CO_2_ emissions and FCR for predicted feeds that minimise CO_2_ footprint (grams CO_2_ per gram growth) based on country-level LCA data for ingredients found in ReCiPe. Results are shown for countries with large Atlantic salmon aquaculture production: Norway, Chile, USA, United Kingdom, and Canada.

The diet recipes from the *in vivo* feed trial [29] consisted of combinations of the same ingredients but with different quality. As shown in Fig. 5, we identified optimal compositions with regards to CO_2_ footprint for the high- and low-quality sets of ingredients. Additional ingredients with available data that could be supplemented to the diets were then identified iteratively with reduction of CO_2_ footprint as objective. The resulting supplemented recipes were similar for Feedtables and ReCiPe, and the optimal mixture in terms of minimal CO_2_ emitted per gram growth was fish meal and fish oil in both cases. Alfalfa protein extract was identified as the best supplement to decrease CO_2_ footprint and also decreased FCR while replacing a substantial amount of fish meal. The best addition for decreasing the CO_2_ footprint further was predicted to be sunflower oil, which increased FCR but decreased the CO_2_ footprint to less than half relative to the initial feed composition. No single ingredient with environmental footprint data could be identified as an addition to further improve this result. The optimal fatty acid composition of the diet with the lowest CO_2_ footprint had the same PUFA content as the original diet, but with a lower *ω*-3-to-*ω*-6 ratio.

**Figure 5:**
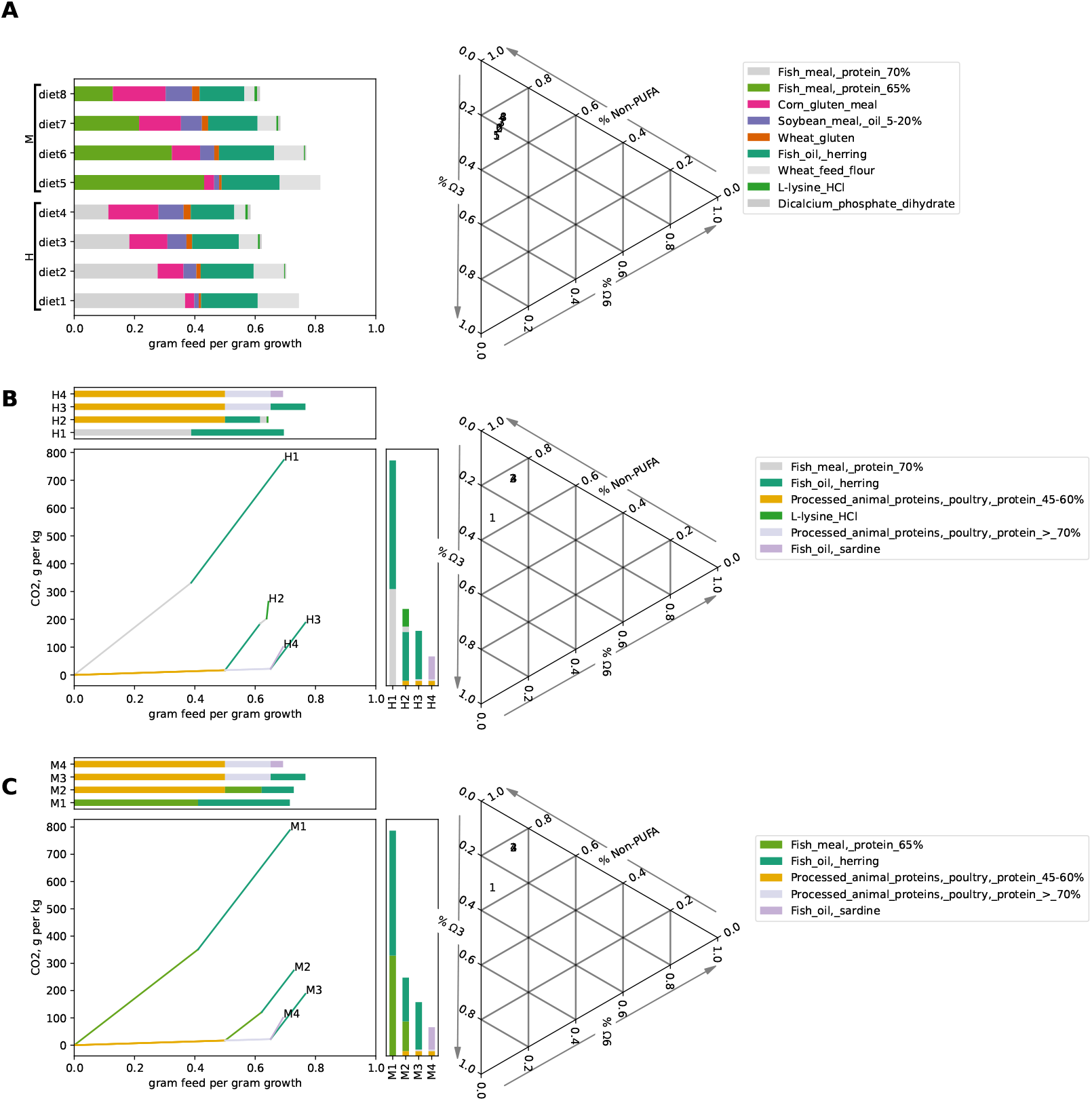
Predicting feed ingredient substitutions that decrease the CO_2_ footprint of an *in vitro* feed trial. (**A**) FCR (grams feed per gram growth) and fatty acid compositions for diets from an *in vitro* feed trial [29] containing fish meal with high (H) or medium (M) quality. The left side shows FCR with optimal compositions sorted from highest to lowest biomass *ω*-3 content. The right side shows the proportion of *ω*-3, *ω*-6, and non-PUFA in the predicted biomass compositions. Diets 1–4 and 5–8 consist of the same ingredients with different ingredient ratios. (**B**) Feed ingredient substitutions that minimise CO_2_ footprint for feeds with high-quality fish meal. Each line represents addition of an ingredient to the diet, ingredients are added in order of increasing contribution to the CO_2_ footprint, and colors identify each ingredient. (**C**) Feed ingredient substitutions that minimise CO_2_ footprint for feeds with medium-quality fish meal.

## 4. Discussion

We have presented SimSaLipiM, a constraint-based metabolic model of Atlantic salmon with flexible and empirically based fatty acid metabolism. The model integrates feed and whole-body composition data, allowing optimization and prediction of key properties such as feed efficiency or fatty acid ratios in salmon biomass. It significantly expands the scope and capabilities of the SALARECON model, specifically when it comes to lipid biosynthesis and degradation, thus allowing detailed simulaton of feed trials *in silico*. If using input data specific to the context of interest, SimSaLipiM can serve as a supplement for *in vivo* feed trials and as a testing ground for hypotheses of novel feed ingredients. The source code is publicly available and can be used to generate models with customised fatty acid metabolism and body composition. It can also easily be used to perform *in silico* feed trials given molecular ingredient compositions and optionally other data such as ingredient prices and environmental footprints.

Fitting the model to data from an *in vivo* feed trial, we predicted efficiency, energy budget, and salmon lipid biomass composition for each tested feed. Diets containing higher proportions of plant-based protein seemed to have higher efficiency, depending on the amount of energy required for maintenance and activity. This variation may be due to the process of mapping ingredients to molecular compositions in Feedtables and the uniform scaling used to match the energy content of the listed diets. A scaling with protein and lipid content taken into account could perhaps have improved the result, but protein and lipid content of each ingredient in the diet would be preferrable. By including the feed conversion ratio in the energy budget, the published energy conversion ratio was approximated, and a detailed energy budget was estimated. The two unique sets of ingredients from the experimental diets were further used to identify optimal recipes based on CO_2_ footprint, and additions that could further improve the efficiency of the diets were also identified. For both unique feed compositions, alfalfa protein concentrate and sunflower oil were identified as beneficial additions to improve climate footprint. However, about half of the *ω*-3 fatty acids in the resulting predicted product was replaced by *ω*-6, highlighting the importance of balancing competing objectives such as efficiency and sustainability.

Using feed ingredient compositions from Feedtables, 58 published fatty acid compositions of salmon that span the space of possible body compositions of fatty acids, and climate footprint as cost in the objective function, a range of optimal feed recipes were predicted, depending on the selected cost data. The CO_2_ footprint listed for a subset of ingredients in Feedtables was used to identify optimal feed recipes based on those ingredients, and the most efficient recipe for a normal feed conversion ratio consisted of processed pig protein, poultry fat, byproducts from biscuit production and linseed. Higher efficiency could be achieved by replacing biscuit byproduct with processed poultry protein and lysine. Mapping ingredents from feedtables to ReCiPe yielded a slightly larger pool of ingredients to choose from, and the most growth-to-emmision-effective feed was predicted to consist of alfalfa protein concentrate and fish oil from capelin for normal FCRs. By using country of origin to filter the climate footprint data in ReCiPe, optimal national feed recipes could be predicted as well, and a feed consisting of Canadian alfalfa protein extract, maize germs, and fish solubles was identified as the optimal feed composition for minimizing CO_2_ footprint per gram fish produced.

SimSaLipiM can be applied in feed design and to gain mechanistic insight into the turnover of compounds and energy in the metabolism of Atlantic salmon. The choice of objective function and constraints is determinative of the result, and with many of these being context-specific, the quality and relevance of input data is key. However, given reasonable compositions of feed ingredients and salmon biomass under given conditions, SimSaLipiM is capable of predicting not only feed efficiency but also the detailed allocation of energy and fatty acids from the feed to the salmon. A feed company or other actors that deal with suppliers of feed ingredients are likely to have access to both LCA data and molecular compositions for their available ingredients, allowing accurate *in silico* analysis of their *in vivo* feed trials.

## 5. Acknowledgements

Funding: This work was supported by the Research Council of Norway grant DigiSal (248792).

## References

[1] The State of World Fisheries and Aquaculture: Meeting the Sustainable Development Goals, Tech. rep., Food and Agriculture Organization of the United Nations (2018).

[2] J. Boissy, J. Aubin, A. Drissi, H. M. Van Der Werf, G. J. Bell, S. J. Kaushik, Environmental impacts of plant-based salmonid diets at feed and farm scales, Aquaculture 321 (1-2) (2011) 61–70.

[3] S. Jennings, G. D. Stentiford, A. M. Leocadio, K. R. Jeffery, J. D. Metcalfe, I. Katsiadaki, N. A. Auchterlonie, S. C. Mangi, J. K. Pinnegar, T. Ellis, E. J. Peeler, T. Luisetti, C. Baker-Austin, M. Brown, T. L. Catchpole, F. J. Clyne, S. R. Dye, N. J. Edmonds, K. Hyder, J. Lee, D. N. Lees, O. C. Morgan, C. M. O’Brien, B. Oidtmann, P. E. Posen, R. Santos, N. G. H. Taylor, A. D. Turner, B. L. Townhill, D. W. Verner-Jeffreys,Aquatic food security: insights into challenges and solutions from an analysis of interactions between fisheries, aquaculture, food safety, human health, fish and human welfare, economy and environment, Fish and Fisheries 17 (4) (2016) 893–938.

[4] M. Bou, G. M. Berge, G. Baeverfjord, T. Sigholt, T.-K. Østbye, Ruyter, Low levels of very-long-chain n-3 PUFA in Atlantic salmon ( Salmo salar) diet reduce fish robustness under challenging conditions in sea cages, Journal of Nutritional Science 6 (2017) e32.

[5] R. K. Saini, Y.-S. Keum, Omega-3 and omega-6 polyunsaturated fatty acids: Dietary sources, metabolism, and significance—A review, Life Sciences 203 (2018) 255–267.

[6] J. O. Agboola, E. M. Chikwati, J. Ø. Hansen, T. M. Kortner, L. T. Mydland, Å. Krogdahl, B. Djordjevic, J. W. Schrama, M. Øverland, A meta-analysis to determine factors associated with the severity of enteritis in Atlantic salmon (Salmo salar) fed soybean meal-based diets, Aquaculture 555 (2022) 738214.

[7] T. S. Aas, T. Åsgård, T. Ytrestøyl, Utilization of feed resources in the production of Atlantic salmon (Salmo salar) in Norway: an update for 2020, Aquaculture Reports 26 (2022) 101316.

[8] M. Zakhartsev, F. Rotnes, M. Gulla, O. Øyås, J. C. J. Van Dam, M. Suarez-Diez, F. Grammes, R. A. Hafþórsson, W. Van Helvoirt, J. J. Koehorst, P. J. Schaap, Y. Jin, L. T. Mydland, A. B. Gjuvsland, S. R. Sandve, V. A. P. Martins Dos Santos, J. O. Vik, SALARECON connects the Atlantic salmon genome to growth and feed efficiency, PLOS Computational Biology 18 (6) (2022) e1010194.

[9] S. Gomez Romero, N. Boyle, Systems biology and metabolic modeling for cultivated meat: A promising approach for cell culture media optimization and cost reduction, Comprehensive Reviews in Food Science and Food Safety 22 (4) (2023) 3422–3443.

[10] B. R. Weston, I. Thiele, A nutrition algorithm to optimize feed and medium composition using genome-scale metabolic models, Metabolic Engineering 76 (2023) 167–178.

[11] H. Molversmyr, O. Øyås, F. Rotnes, J. O. Vik, Extracting functionally accurate context-specific models of Atlantic salmon metabolism, npj Systems Biology and Applications 9 (1) (2023) 19.

[12] A. M. Feist, B. O. Palsson, The biomass objective function, Current Opinion in Microbiology 13 (3) (2010) 344–349.

[13] C. Schulz, T. Kumelj, E. Karlsen, E. Almaas, Genome-scale metabolic modelling when changes in environmental conditions affect biomass composition, PLoS Computational Biology 17 (5) (2021) e1008528.

[14] V. Simensen, C. Schulz, E. Karlsen, S. Bråtelund, I. Burgos, L. B. Thorfinnsdottir, L. García-Calvo, P. Bruheim, E. Almaas, Quantification of macromolecular biomass composition for constraint-based metabolic modeling, bioRxiv (2021) 2021–08.

[15] J. G. Bell, I. Ashton, C. J. Secombes, B. R. Weitzel, J. R. Dick, J. R. Sargent, Dietary lipid affects phospholipid fatty acid compositions, eicosanoid production and immune function in Atlantic salmon (Salmo salar), Prostaglandins, Leukotrienes and Essential Fatty Acids 54 (3) (1996) 173–182.

[16] D. R. Tocher, J. G. Bell, J. R. Dick, J. R. Sargent, Fatty acyl desaturation in isolated hepatocytes from Atlantic salmon (Salmo salar): Stimulation by dietary borage oil containing gamma-linolenic acid, Lipids 32 (12) (1997) 1237–1247.

[17] K. Tummler, E. Klipp, The discrepancy between data for and expectations on metabolic models: How to match experiments and computational efforts to arrive at quantitative predictions?, Current Opinion in Systems Biology 8 (2018) 1–6.

[18] B. J. Sánchez, F. Li, E. J. Kerkhoven, J. Nielsen, SLIMEr: probing flexibility of lipid metabolism in yeast with an improved constraintbased modeling framework, BMC Systems Biology 13 (1) (2019) 4.

[19] H. Lu, F. Li, B. J. Sánchez, Z. Zhu, G. Li, I. Domenzain, S. Marcišauskas, P. M. Anton, D. Lappa, C. Lieven, et al., A consensus S. cerevisiae metabolic model Yeast8 and its ecosystem for comprehensively probing cellular metabolism, Nature Communications 10 (1) (2019) 3586.

[20] E. Brunk, S. Sahoo, D. C. Zielinski, A. Altunkaya, A. Dräger, N. Mih, F. Gatto, A. Nilsson, G. A. Preciat Gonzalez, M. K. Aurich, et al., Recon3D enables a three-dimensional view of gene variation in human metabolism, Nature Biotechnology 36 (3) (2018) 272–281.

[21] J. L. Robinson, P. Kocabaş, H. Wang, P.-E. Cholley, D. Cook, A. Nilsson, M. Anton, R. Ferreira, I. Domenzain, V. Billa, et al., An atlas of human metabolism, Science Signaling 13 (624) (2020) eaaz1482.

[22] H. Wang, J. L. Robinson, P. Kocabas, J. Gustafsson, M. Anton, P.-E. Cholley, S. Huang, J. Gobom, T. Svensson, M. Uhlen, et al., Genomescale metabolic network reconstruction of model animals as a platform for translational research, Proceedings of the National Academy of Sciences 118 (30) (2021) e2102344118.

[23] D. R. Tocher, Metabolism and functions of lipids and fatty acids in teleost fish, Reviews in Fisheries Science 11 (2) (2003) 107–184.

[24] L. van Steijn, F. J. Verbeek, H. P. Spaink, R. M. Merks, Predicting metabolism from gene expression in an improved whole-genome metabolic network model of Danio rerio, Zebrafish 16 (4) (2019) 348–362.

[25] J. D. Orth, I. Thiele, B. Ø. Palsson, What is flux balance analysis?, Nature Biotechnology 28 (3) (2010) 245–248.

[26] INRAE-CIRAD-AFZ Feed tables, https://www.feedtables.com/.

[27] T. S. Mock, D. S. Francis, D. W. Drumm, V. L. Versace, B. D. Glencross, R. P. Smullen, M. K. Jago, G. M. Turchini, A systematic review and analysis of long-term growth trials on the effect of diet on omega-3 fatty acid levels in the fillet tissue of post-smolt Atlantic salmon, Aquaculture 516 (2020) 734643.

[28] C. J. Norsigian, N. Pusarla, J. L. McConn, J. T. Yurkovich, A. Dräger, B. O. Palsson, Z. King, Bigg models 2020: multi-strain genome-scale models and expansion across the phylogenetic tree, Nucleic acids research 48 (D1) (2020) D402–D406.

[29] H. Mundheim, A. Aksnes, B. Hope Growth, feed efficiency and digestibility in salmon (Salmo salar L.) fed different dietary proportions of vegetable protein sources in combination with two fish meal qualities, Aquaculture 237 (1) (2004) 315–331.

[30] N. E. Lewis, K. K. Hixson, T. M. Conrad, J. A. Lerman, P. Charusanti, A. D. Polpitiya, J. N. Adkins, G. Schramm, S. O. Purvine, D. Lopez-Ferrer, et al., Omic data from evolved E. coli are consistent with computed optimal growth from genome-scale models, Molecular Systems Biology 6 (1) (2010) 390.

[31] Factors affecting energy and protein requirements, https://www.fao.org/4/aa040e/AA040E08.htm.

[32] Global Feed LCA Institute (GFLI) database, https://globalfeedlca.org/gfli-database/.

[33] A. Richelle, B. P. Kellman, A. T. Wenzel, A. W. Chiang, T. Reagan, J. M. Gutierrez, C. Joshi, S. Li, J. K. Liu, H. Masson, et al., Model-based assessment of mammalian cell metabolic functionalities using omics data, Cell Reports Methods 1 (3) (2021).

